# Describing the Local Structure of Sequence Graphs

**DOI:** 10.1101/125542

**Authors:** Yohei Rosen, Jordan Eizenga, Benedict Paten

## Abstract

Analysis of genetic variation using graph structures is an emerging paradigm of genomics. However, defining genetic sites on sequence graphs remains an open problem. Paten’s invention of the *ultrabubble* and *snarl*, special subgraphs of sequence graphs which can identified with efficient algorithms, represents important first step to segregating graphs into genetic sites. We extend the theory of ultrabubbles to a special subclass where every detail of the ultrabubble can be described in a series and parallel arrangement of genetic sites. We furthermore introduce the concept of *bundle* structures, which allows us to recognize the graph motifs created by additional combinations of variation in the graph, including but not limited to runs of abutting single nucleotide variants. We demonstrate linear-time identification of bundles in a bidirected graph. These two advances build on initial work on ultrabubbles in bidirected graphs, and define a more granular concept of genetic site.

## 1 Background

The concept of the genetic site underpins both classical genetics and modern genomics. From a biological perspective, a site is a position at which mutations have occurred in different samples’ histories, leading to genetic variation. From an engineering perspective, a site is a subgraph with left and right endpoints where traversals by paths correspond to alleles. This is useful for indexing and querying variants in paths and for describing variants in a consistent and granular manner.

Against a linear reference, it is trivial to define sites, provided that we disallow variants spanning overlapping positions. This is clearly demonstrated by VCF structure [1]. VCF sites, consisting of any number of possible alleles, are identified by their endpoints with respect to the linear reference.

If we wish to analyze a set of variants containing structural variation, highly divergent sequences or nonlinear references structures, then a linear reference with only non-overlapping variants is no longer a sufficient model. Datasets with one or more of these properties are becoming more common [2, 3], and sequence graphs [4] have been developed as a method of representing them. However, defining sites on graphs is considerably more difficult than on linear reference structures and the creation of methods to fully decompose sequence graphs into sites remains an unsolved problem.

## 2 The Challenges of Defining Sites on Graphs

On a graph-based reference, the linear reference definition of a site as a position along the reference and a set of alleles fails to work for several reasons:

1. 1.Sequences which are at the same location in linear position may not have comparable contexts. This is a consequence of having variants which cannot be represented as edits to the linear reference but rather as edits to another variant. We illustrate this with an example from *1000 Genomes* polymorphism data, visualized using Sequence Tube Maps [5].
2. Elements of sequence may not be linearly ordered. Parallel structure of the graph (3.) is one sort of non-linearity. Graphs also allow repetitive, inverted or transposed elements of sequence. These all prevent linear ordering.
3. The positions spanned by different elements of variation may partially overlap. Therefore, multiple mutually exclusive segments of sequence in a region of the graph cannot be considered to be alternates to each other at a well-defined position without having to include extraneous sequence that is shared between some but not all of the “alleles.”

We can expect that the density of these graph structures will increase with increasing population sizes included in datasets.

Our aim will be to recognize and fully decompose subgraphs resembling Example 1 into a notion of site, and isolate these from elements of the graph resembling Examples 2 and 3.

## 3 Mathematical Background

### 3.1 Directed and Bidirected Sequence Graphs

The graphs used to represent genetic information consist of labelled nodes and edges. Nodes are labelled with sequence fragments. Edges form paths whose labels spell out allowed sequences. Two types of graph are used.

The more simple type is the directed graph. A directed graph (or “digraph”) *G* consists of a set *V* of nodes and a set *E* of directed edges. A directed edge is an ordered tuple (*x, y*), consisting of a *head x* ∈ *V* and *tail y* ∈ *V*. A directed path is a sequence of nodes joined by edges, followed head to tail. *G* is a *directed acyclic graph* (DAG) if it admits no directed path which revisits any node.

A bidirected graph *G* [6] consists of a set *V* of vertices and a set *E* of edges. Each vertex *υ* ∈*V* consists of a pair of *node-sides* {*υ_left_, υ_right_*} and each edge is an unordered tuple of node-sides. Bidirected graphs have the advantage of being able to represent inversion events.

We write *N* for the set of node-sides in the bidirected graph *G*. The *opposite* 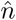 of a node-side *n* is the other node-side at the same vertex as *n*.

A sequence *p* = *x*_1_, *x*_2_, …, *x_k_* of node-sides is a *path* if ∀*x_i_*,

1. if 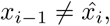, then 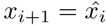
2. if 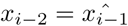, then {*x*_*i*−1_, *x_i_*} ∈ *E*
3. any contiguous subsequence of *p* consisting of a node-side *x* alternating with its opposite 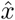 must either be even-numbered in length or must be a prefix or suffix of *p*

Informally, this means that in a path, consecutive pairs forming edges alternate with pairs of opposite node-sides or, equivalently, that paths visit both node-sides of the vertices they pass through. They can however begin or on an isolated node-side.

A bidirected graph *G* is *cyclic* if it admits a path visiting a node-side twice. Therefore the self-incident hairpin motif (below, right) is considered a cycle. A bidirected graph *G* is *properly cyclic* if it admits a path which visits a pair 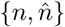 twice in the same order.

Some publications refer to biedged graphs. These are {*black, grey*}-edge-colored undirected graphs, where every node is paired with precisely one other by sharing a grey edge and paths in the graph must alternate between traversing black and grey edges. Paten elaborates on this construction in [7] and shows that it is equivalent to a bidirected graph. We will restrict our language to that of bidirected graphs, recognizing that these are equivalent to biedged graphs.

Acyclic bidirected graphs are structurally equivalent to directed graphs in that

#### Lemma 1.

*If G is a bidirected acyclic graph, there exists an isomorphic directed acyclic graph D*(*G*).

*Proof. See [7]*

### 3.2 Bubbles, Superbubbles, Ultrabubbles and Snarls

The first use of local graph structure to identify variation was the detection of *bubbles* [8] in order to detect and remove sequencing errors from assembly graphs. Their bubble is the graph motif consisting of two paths which share a source and a sink but are disjoint between.

The general concept of bubbles was extended by Onodera et al, who defined superbubbles in directed graphs [9]. Brankovic demonstrates an 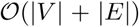 algorithm to identify them [10], building off work of Sung [11].

We restate the Onodera definition, modified slightly as to be subgraph-centric rather than boundary-centric: A subgraph *S* ⊆ *G* of a directed graph is a *super-bubble with boundaries* (*s, t*) if

1. (reachability) *t* is reachable from *s* by a directed path in *S*
2. (matching) the set of vertices reachable from *s* without passing through *t* is equal to the set of vertices from which *t* is reachable without passing through *s*, and both are equal to *S*
3. (acyclicity) *S* is acyclic
4. (minimality) there exists no *t*′ ∈ *S* such that boundaries (*s, t*′) fulfil 1,2 and 3.There exists no *s*′ ∈ *S* such that (*s*′, *t*) fulfil 1,2 and 3.

To motivate our definition of a superbubble equivalent on bidirected graphs, we prove some consequences of the matching property.

#### Proposition 2.

*Let S ⊆ *G* be a subgraph of a directed graph. If S possesses the matching property relative to a pair* (*s,t*), *then it possesses the following three properties:*

1. *(2-node separability) Deletion of all incoming edges of s and all outgoing edges of t disconnects S from the remainder of the graph*.
2. *(tiplessness) There exist no node n* ∈ *S*\{*s, t*} *such that n has either only incoming or outgoing edges*.
3. *S is weakly connected*

*Proof*. (*matching* ⇒ *separability*) *Suppose* Ǝ *x* ∉ *S, y* ∈ *S*\{*s, t*} *such that there exists either an edge x* → *y or an edge y* → *x. Suppose wlog that* ∃ *an edge x* → *y. By matching, there exists a path y* → · · · → *t without passing through s. We can then construct the path x → y* → · · · → *t which does not pass through s. But by matching this implies that x* ∈ *S, which leads to a contradiction*.

The converse need not be true on directed graphs^3^. We define two structures on bidirected graphs. The first is the ultrabubble, which given Proposition 2, can be thought of as an analogue to a superbubble. The second, the snarl, is a more general object which preserves the property of 2-node separability from the larger graph without having strong guarantees on its internal structure. The following definitions are due to Paten [7]:

A connected subgraph *S* ⊆ *G* of a bidirected graph *G* is a *snarl* (*S, s, t*) with boundaries (*s, t*), if

1. 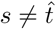
2. (2-node separability) every path between a pair of node-sides in *x* ∈ *S, y* ∈ *G*\*S* contains either 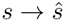 or 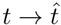 a subpath.
3. (minimality) there exists no *t*′∈ *S* such that boundaries (*s, t*′) fulfil 1 and 2.There exists no *s*′∈ *S* such that (*s*′, *t*) fulfil 1 and 2 The class of *ultrabubbles* is the subclass of snarls (*S, s, t*) furthermore fulfilling
4. 4. S is acyclic
5. S contains no tips — vertices having one node-side involved in no edges Three examples of ultrabubbles are shown below.

The following is important property of snarls.

#### Proposition 3 (Non-overlapping property).

*If two distinct snarls share a vertex (node-side pair) then either they share a boundary node or one snarl is included in the other’s interior*.

*Proof. Let S be a snarl with boundaries s, t. Let T be another snarl, with bound-aries u, υ*. *Suppose that u* ∈ *S*\{*s,t*} but *υ* ∉ *S*, and *s* ∉ *T*.

*Consider the set *S* ∩ *T*. It is nonempty since it contains *u*. Let *x* ∈ *S* ∩ *T*. Let *y* ∉ *S* ∩ *T*. Suppose that there exists a path *p* = *x* ↔ · · · ↔ *y* which neither passes through u nor t*.

*Since y* ∉ *S* ∩ *T, either y* ∉ *S or y* ∉ *T. Wlog, assume y* ∉ *T. Then due to the separability of T, since the path p does not pass through u, it must pass through v before leaving T to visit y. But v* ∉ *S so p must also pass through s before leaving S to visit v since it does not pass through t. But it must pass through v before leaving T to visit s, which leads to an impossible sequence of events. Therefore any path x ↔ · · · ↔ y* for x *∈* S *∩*T, y∉ S*∩ T must pass through either u or t. This contradicts the minimality of both S and T*.

This non-overlapping property is also a nesting property. Observe that, due to Proposition 3, the relation *U* ≤ *υ* on snarls *U, V* defined such that *U* ≤ *V* if *U* is entirely contained in *V* has the property that if *U* ≤ *V* and *U* ≤ *W*, then either *V* ≤ *W* or *W* ≤ *V*. Therefore the partial order on the snarls of *G* defined by the relation ≤ will always be equivalent to a tree diagram. A *bottom level* snarl is one which forms a leaf node of this tree.

The equivalent of Proposition 3 for superbubbles was stated without proof by Onodera in [9]. Our proof also constitutes a proof of the statement for super-bubbles, due to the following proposition, proven by Paten in [7]:

#### Proposition 4.

*Every superbubble in a directed graph corresponds to an ultrabubble in the equivalent (see Lemma 1) bidirected graph*.

Identifying all superbubbles in a directed graph or all snarls in a bidirected graph introduces a method of compartmentalizing a graph into partitions whose contents are all in some sense at the same position in the graph, and for which the possible internal paths are independent of what path they continue on beyond their boundaries. We will use this concept to define sites for certain specialized classes of graphs.

## 4 Graphs which are Decomposable into Nested Simple Sites

We will extend the theory of ultrabubbles to a theory of nested sites where the structure of certain graphs can be fully described in terms of combinations of linear orderign and ultrabubble nesting relationships. This is important for

1. Identifying nested variation
2. Indexing traversals

### 4.1 Traversals and Subpaths

An (*s, t*)-*traversal* of *S* is a path in *S* beginning with *s* and ending with *t*. An (*s, s*)*-traversal* and a (*t*,*t*)-*traversal* are analogously defined. Presence of an (*s, s*)- or (*t, t*)-traversal implies cyclicity. Two traversals of a snarl are *disjoint* if they are disjoint on *S*\{*s*,*t*}.

Paten’s [7] snarls and ultrabubbles are 2-node separable subgraphs whose paired boundary nodes isolate their traversals from the larger graph. We can state this with more mathematical rigor:

*Claim*. Consider a snarl (*S, s*,*t*) in a bidirected graph *G*. The set of all paths in *G* which contain a single (*s, t*)-traversal as contiguous a subpath is isomorphic to the set-theoretic product *P*(*s*) × *Trav*(*s, t*) × *P*(*t*) consisting of the three sets

1. *P*(*s*) := {paths in *G*\*S* terminating in 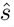}
2. *Trav*(*s*,*t*) := {(*s, t*)-traversals of *S*}
3. *P*(*t*) := {paths in *G*\*S* beginning 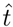}

The isomorphism is the function mapping *p*_1_ ∈ *P*(*s*), *p*_2_ ∈ *Trav*(*s*,*t*), *p*_3_ ∈ *P*(*t*) to their concatenation *p*_1_*p*_2_*p*_3_.

This property is important because it allows us to express the set of all haplotypes traversing a given linear sequence of snarls in terms of combinations of alleles for which we do not need to check if certain combinations are valid.

### 4.2 Simple Bubbles and Nested Simple Bubbles

#### Definition 5.

*An ultrabubble* (*S, s, t*) *is a simple bubble if all* (*s, t*)*-traversals are disjoint*.

Simple bubbles are structurally equivalent to (multiallelic) sites consisting of disjoint substitutions, insertions or deletions, with all alleles spanning the same boundaries.

Proposition 7 below demonstrates that we can identify simple bubbles in 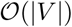 time given that we have found all snarl boundaries. Paten has shown [7] that identification of snarl boundaries is achieved in 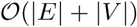 time. To find the ultrabubbles among these, note that checking for acyclicity is 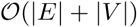 on account of the unbranching nature of these snarls’ interiors.

Given a node-side *n*, write *Nb*(*n*) for the set of all neighbors of *n*. Note that *a* ∈ *Nb*(*b*) ⇔ *b* ∈ *Nb*(*a*).

#### Lemma 6 (Nodes in an ultrabubble are orientable with respect to the ultrabubble boundaries).

*Given an ultrabubble* (*S, s, t*) *and given n* ∈ *S\*{*s, t*}, *consider the set T of all* (*s,t*)-*traversals of S passing through n. Then either*

1. ∀_*p*_ *∈ T*, an element of *Nb*(*n*) *precedes n in p*
2. ∀_*p*_ *∈ T, an element of Nb*(*n*) *follows n in p*

*In the former case we call n s-sided, otherwise we call it t-sided*.

*Proof. This is a corollary to Lemma 1*.

#### Proposition 7 (Simple bubbles have unbranching interiors).

*Let* (*S, s, t*) *be an ultrabubble. Then all traversals are disjoint iff every interior node-side has precisely one neighbor*.

*Proof*. (⇒) *Suppose that all* (*s, t*)-*traversals of S are disjoint. Suppose* Ǝ *a node-side n* ∈ *S*\{*s, t*}* with multiple neighbors*.

*Since n is orientable with respect to* (*s, t*), *suppose, without loss of generality, that it is s-sided. Then there exist distinct paths from s to n passing through each of its neighbors. Continuing these with a path from n^opp^ to t produces two nondisjoint traversals of S*.

(⇐) *Suppose that every interior node-side has precisely one neighbor. Suppose that there exist two distinct nondisjoint traversals of S. For no node-side to have multiple neighbors, they must coincide at every node-side, contradicting the assumption that they are not the same traversal*.

We seek to extend this simple property to more complex graph structures. We will take advantage of the nesting of nondisjoint ultrabubbles proven in Proposition 3 to define another structure in which nondisjoint traversals are easily indexed.

#### Definition 8.

*An ultrabubble* (*S, s, t*) ⊆ *G is decomposable into nested simple sites if either*:

1. *S is a simple bubble*
2. *if, for every ultrabubble S′ contained in the interior of S, you replace the ultrabubble with a single edge s — t whenever S′ is decomposable into simple sites, then S becomes a simple bubble*

The following figure demonstrates decomposability into nested simple sites.

#### Proposition 9.

*If an ultrabubble* (*U, s, t*) *is decomposable into nested simple sites, then the complete node sequence of any* (*s, t*)-*traversal can be determined only by specifying the path it takes inside those nested ultrabubbles within which the traversal does not visit any further nested ultrabubble*.

*Proof. Let p be a* (*s, t*)*-traversal of an ultrabubble U which is decomposable into nested simple sites. Let V be a nested ultrabubble inside U. If p traverses, V, write p|_V_ for the traversal p restricted to V*

*Suppose that t*|_*V*_ *intersects no nested ultrabubbles within V. Then t*|_*V*_ *is disjoint of all other traversals within V due to U begin decomposable into nested simple sites. Therefore specifying any node of t|V uniquely identifies it*.

*Suppose that t*|_*V*_ *intersects some set of ultrabubbles nested within V. Since U is decomposable into nested simple sites, the nodes of t*|_*V*_ *must be linear and disjoint of all other paths if we replace all ultrabubbles nested in V with edges joining their boundaries. Therefore specifying which ultrabubbles are crossed uniquely determines the nodes included in t*|_*V*_ *which lie outside of the nested ultrabubbles in V*.

*The statement of the proposition follows from the two arguments above by induction*.

#### Proposition 10.

*An ultrabubble is decomposable into nested simple sites iff every node side is either the interior ultrabubble boundary or has precisely one neighbor*.

*Proof. This can be established using Proposition 7*.

This property allows 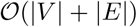evaluation of whether a graph is decomposable into nested simple sites, by arguments analogous to those for simple bubbles.

### 4.3 A Partial Taxonomy of Graph Notifs which do not Admit Decomposition into Sites

In section 4.3, we will show that we can decompose a graph into nested simple sites as defined in the previous section if it lacks a certain forbidden motif. We will begin with examples of three graph motifs, and the biological events which might produce them.

We describe some graph features which prevent decomposition into nested sites below, and the sets of mutations which might have produced them.

1. Two (or more) substitutions or deletions against a linear sequence which overlap, but not completely.
2. A substitution (or deletion) which spans elements of sequence on the interior of two disjoint ultrabubbles. Addition of such an edge joining two ultrabubbles which were decomposable into nested simple sites will consolidate the two into a single ultrabubble which is not decomposable into nested simple sites.
3. Two SNVs or other simple elements of variation at adjacent positions. This will be the focus of our Section 5.

### 4.4 The Relationship Between Nested Simple Sites and Series Parallel Graphs

The structure of ultrabubbles decomposable into nested simple sites, and their tree representation (see Fig 7) might be familiar to the graph theorist familiar with series-parallel digraphs. The fact that the digraphs equivalent to ultrabubbles form a subclass of the two-terminal series-parallel digraphs is interesting due to the computational properties of the latter class of graphs.

**Fig.1.**
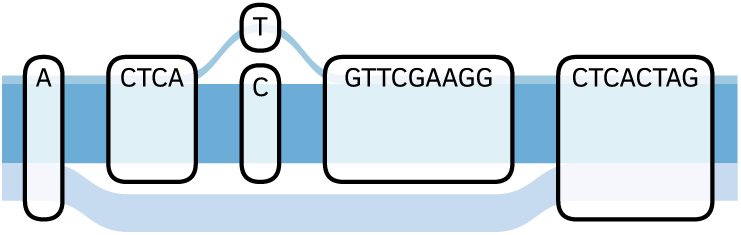
The context of the single nucleotide variant shown does not exist in all variants spanning its linear position

**Fig. 2.**
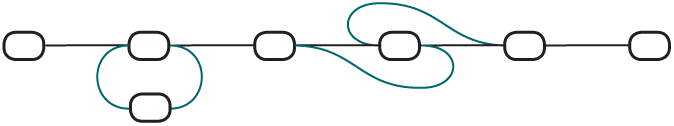
A cycle and an inversion in a graph

**Fig. 3.**
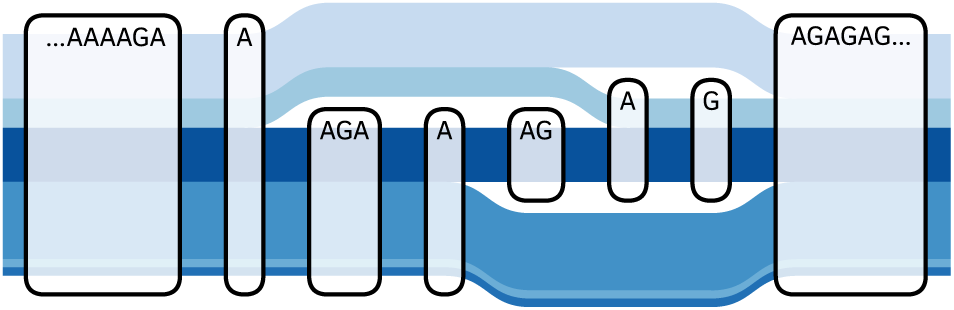
Overlapping deletions, from 1000 Genomes polymorphism data

**Fig. 4.**
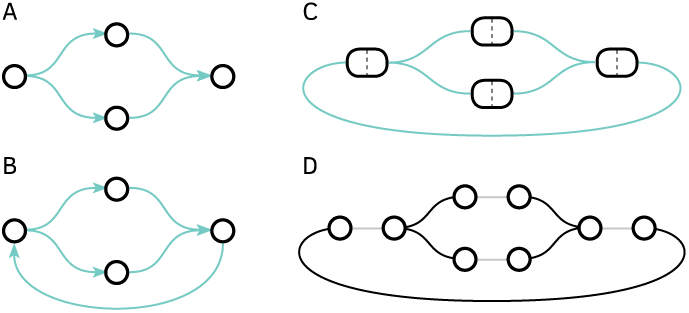
(A) A directed acyclic graph (B) A cyclic directed graph (C) Graph B represented as a bidirected graph. This cycle is proper. (D) Graph C represented as a biedged graph

**Fig. 5.**
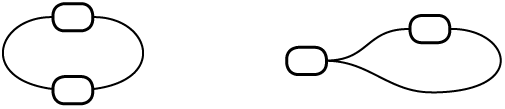
(Left) A properly cyclic graph. (Right) The self-incident hairpin motif of a cyclic but not properly cyclic graph

**Fig. 6.**
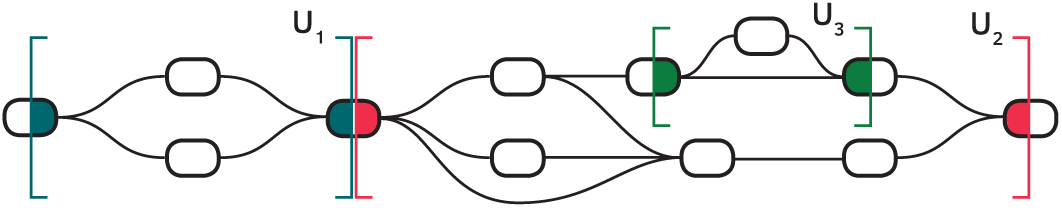
Three ultrabubbles, boundaries colored blue, pink and green. These illustrate the non-overlapping property

**Fig. 7.**
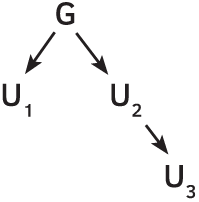
The nesting tree diagram of the ultrabubbles from the previous figure. *U*_1_ and *U*_3_ are bottom-level

#### Definition 11.

*A directed graph G is two-terminal series parallel (*TTSP*) with source s and sink t if either*

1. *G is the two-element graph with a single directed edge s *→* t*
2. *There exist TTSP graphs G*_1_, *G*_2_ *with sources s*_1_, *s*_2_ and sinks *t*_1_, *t*_2_ *such that G is formed from G*_1_, *G*_2_ *by identification of s*_1_ *with s*_2_ *as s and identification of t*_1_ with *t*_2_ as *t(Parallel addition)*
3. *There exist TTSP graphs G*_1_, *G*_2_ with sources *s*_1_, *s*_2_ and sinks *t*_1_, *t*_2_ *such that G is formed, from G*_1_, *G*_2_ *by identification of *t*_1_ with *s*_2_ (Series addition)*

Two terminal series parallel digraphs have a useful forbidden subgraph characterization.

#### Proposition 12 (From [12]).

*A directed graph G is two terminal series parallel if and only if it contains no subgraph homeomorphic to the graph W shown below Proof: Refer to Valdes [12] and Duffin [13]*

#### Proposition 13.

*If an ultrabubble* (*U, s, t*) *is decomposable into nested simple sites, then the equivalent directed graph is TTSP with source s and sink t*.

*Proof. Suppose that the directed graph D*(*U*) *equivalent to U (which exists by Lemma 1) contains a subgraph homeomorphic to W. Then there must be a node-side u in U with two neighbours a*_1_, *a*_2_ *which are the beginnings of disjoint paths p*_1_, *p*_2_ *ending on node-sides b*_1_, *b*_2_ *which are neighbours of a node-side v. By Proposition 10, u and v must be ultrabubble boundaries. Since p*_1_, *p*_2_ *are disjoint, u and v must be opposing boundaries of the same ultrabubble. But the presence of a subgraph homeomorphic to W also implies that there exists a pair q*_1_, *q*_2_ *of disjoint paths, one from a node x to* 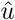 *and the other from x to v, both not passing through* 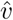. *But this is not possible since it would contradict 2-node separability of* (*u, υ*).

We highlight the middle “Z-arm” of the *W*-motif in our first two examples of ultrabubbles which are not decomposable into nested simple sites.

## 5 Abutting Variants

We wish to decompose the graph structure of sets of variants lying at adjacent positions such that there is no conserved sequence between them able to form an ultrabubble boundary. We will define a graph motif called the *balanced recombination bundle* which corresponds this graph structure, and can be rapidly detected.

We observe examples abutting single nucleotide variants (SNVs) in the 1000 Genomes polymorphism data. It is a reasonable hypothesis that these should become more common as the population sizes of sequencing datasets increases, since, statistically, the distribution of variation across the genome should grow less sparse as the population increases.

### 5.1 Bundles

#### Definition 14.

*An internal chain n*_1_ *→ n*_2_ → · · · → *n_k_* is a sequence of node-sides such that ∀*i*, 2 ≤ *i* ≤ *k, n_i_* ∈ *Nb*(*n*_*i*−1_).

#### Definition 15.

*We say that a tuple* (*L, R*) *of sets of node-sides is a bundle if*

1. *(Matching)* ∀*ℓ* ∈ *L, Nb*(*ℓ*) ⊆ *R and Nb*(*ℓ*) ≠ ∅; ⊆ _r_ ∈ *R Nb*(*r*)⊆ *L and Nb*(*r*) ≠ ∅
2. *(Connectedness)* ∀*ℓ* ∈ *L, r* ∈ *R, there exists an internal chain ℓ* → *r*_1_ → *ℓ*_1_ → · · · → *r_k_ →ℓ_k_ →* r such that *∀i*, 1 ≤ i ≤ *k, r_i_* ∈ *R and ℓ_i_* ∈ *L*

#### Definition 16.

*We say that a tuple (*L, R*) of sets of node-sides is a balanced recombination bundle (R-bundle for short) if*

1. *(Complete matching)* ∀*ℓ ∈ L, Nb*(*ℓ*) = *R and ∀r* ∈ *R, Nb*(*r*) = *L*
2. *(Acyclicity) L* ∩ *R* = ∅

#### Lemma 17.

*A balanced recombination bundle is a bundle*.

*Proof. Complete matching ⇒ matching*.

*Complete matching *⇒* connectedness by the chain ℓ *→*r for all ℓ *∈ L, r ∈* R*

#### Definition 18.

*An unbalanced bundle is a bundle which is not a balanced recombination bundle. An unbalanced bundle is acyclic if L*∩* R* = ∅

#### Definition 19.

*We say that two bundles* (*L*_1_, *R*_1_), (*L*_2_, *R*_2_) *are isomorphic if either L*_1_ = *L*_2_ *and R*_1_ = *R*_2_ or *L*_1_ = *R*_2_ and *R*_1_ = *L*_2_.

We will describe a 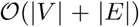algorithm to detect and categorize bundles exhaustively for all node-sides in a bidirected graph. To establish the validity of this algorithm, we need several preliminary results:

#### Lemma 20.

*Every q* ∈ *N is either a tip or an element of a bundle*.

*Proof. Suppose that q is not a tip. Define a function W that maps a tuple* (*L, R*) *of nonempty sets of node-sides to a tuple W*(*L*), *W*(*R*) *where*

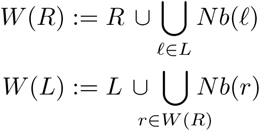

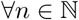 define

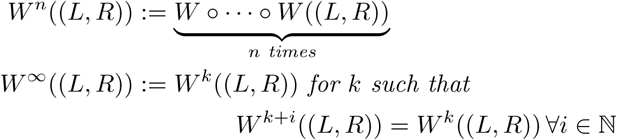

*W^∞^ exists since W^n^ is nondecreasing with respect to set inclusion and our graphs are finite. Now define*, 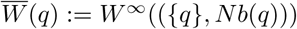, noting that Nb(*q*)≠ ∅ *since*{*q*} *is not a tip. Let us write *L_w_*∞ and R_w_*∞ *for the respective elements of. 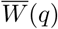. We claim that 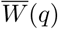 *is a bundle**.

*Proof of matching: let ℓ* ∈ *L_W*∞*,_* r *∈* RW*∞. By construction of W*,

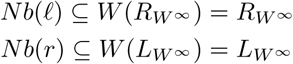

*Proof of connectedness*: *let ℓ ∈*L_W_∞, *r* ∈ *R_W_*∞. *We will show that for any r* ∈ *R_W_*∞, ∃ *an internal chain q → r_1_ →* l*_1_ →· · ·→*r_k_ *→* i_k_ *→ r such that* ∀_*i*_, 1 ≤ *i ≤ k, r_i_ ∈ R_w_*∞ *and* ℓ_*i*_ *∈* L_W_*∞*.

*Suppose that r ∈* Nb(*q), then we are done. Otherwise, since r *∈ Rw∞*, there exists some minimal 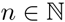*such that r* ∈ the R-set RW ^n^ of some W *^n^(({q}*, Nb(*q*)))*. *It is straightforward to see that we can then construct an internal chain q → r_0_ →* ℓ*_1_ → r*_1_ → … ℓ*_n−1_ →* r such that ∀i*, 1 ≤* i *≤* n *-* 1, r_i_ *∈* R_W_^i^, ℓ_i_ *∈* L_W_^i^. *By an analogous argument, we can do the same for an internal chain ℓ →· · · → r*′ *for some r*′ ∈ *Nb*(*q*). *Concatenation of the first chain with the reverse of the second gives our chain* ℓ → · · · → *r, proving connectedness*.

#### Proposition 21.

*If q *∈* L for a bundle (*L, R*), *then* 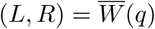*

*Proof. Suppose that 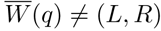. *Then either L ≠ L_W_∞ or R* ≠ *R_W_*∞ First, suppose the latter. Suppose that *Ǝr ∈* R such that r∉RW*∞*. Since (*L, R*) is a bundle, we know that there is an internal chain q*→* r*_0_ →* ℓ*_1_ →* r*_1_ → ··· →* r_k_ *→* i_k_ *→* r with all r_i_ *∈* R, i_i_ *€* L. But, using the same shorthand as before, it is also evident that r_i_ *∈* R,W^i^, i_i_ *∈* LW ^i^ ∀i*, 1 ≤* i *≤* k. But since i_k_ Nb(*r*), we can deduce that r ∈* RW*^k+1^, which leads to a contradiction since r ∈ R*_*W*_∞.

*Suppose otherwise that* ∃*r* ∈ *R_W_∞ such that r ∉R. Consider an internal chain c = q* → *r*_0_ → ℓ_1_ *→ r*_1_ → · · · → *r_k_ →* ℓ_*k*_*→ r fulfilling the conditions needed to prove connectedness of*. 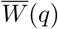. Note that q *∈* L and by matching r_0_ *∈ Nb*(*q*). But r *∈ R, which leads to a contradiction since it means that there must exist two consecutive members somewhere in the chain c which cannot be neighbors*.

We say that a node-side *n* is *involved in* a bundle (*L*,*R*) if *n ∈ L* or *n ∈* R.

#### Corollary 22 (To Proposition 21).

*Every non-tip node-side is involved in precisely one bundle*.

### 5.2 An Algorithm for Bundle-Finding

The diagram in Fig 16 demonstrates our algorithm for finding the balanced recombination bundle containing a query node-side *q* if it is contained in one, and discovering that it is not if it is not. The is written in pseudocode below, with an illustration following.

**Fig. 8.**
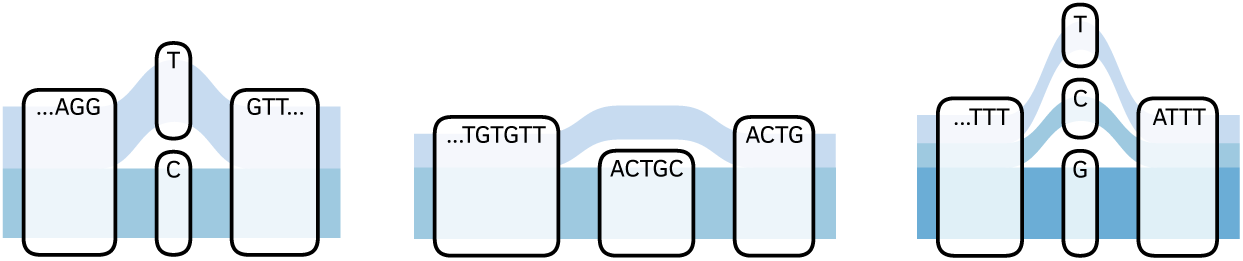
Three examples of simple bubbles from the 1000 Genomes graph

**Fig. 9.**
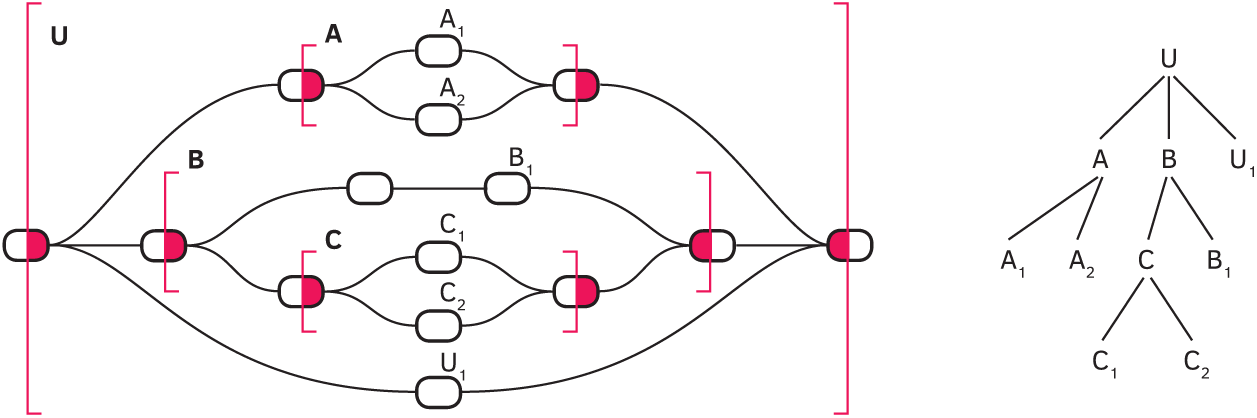
Left: A nesting of four ultrabubbles. Right: The tree structure to index traversals of *U* implied by Proposition 9

**Fig. 10.**
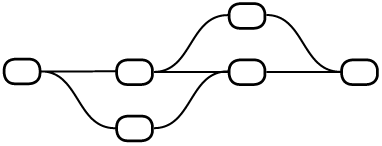
Overlapping substitutions (or deletions)

**Fig. 11.**
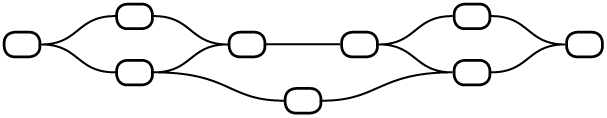
An edge crossing bubble boundaries.

**Fig. 12.**
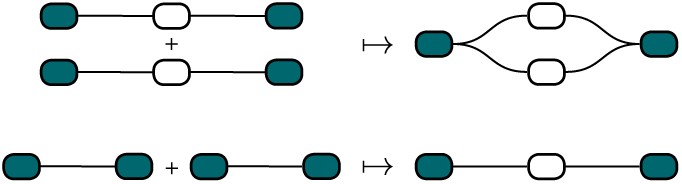
Top: parallel addition. Bottom: series addition

**Fig. 13.**
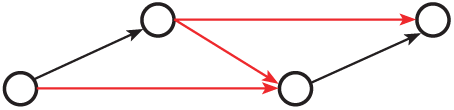
The W motif

**Fig. 14.**
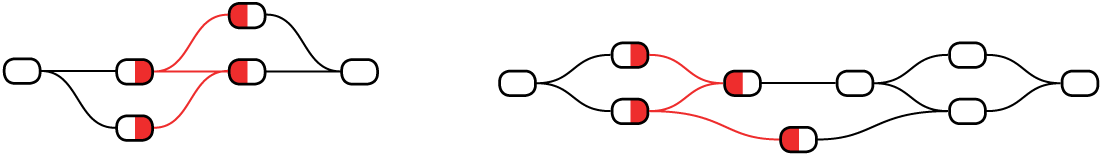
Portions of the ultrabubbles 1. and 2. of section 4.2, showing the nodes which project to the forbidden subgraph *W*

**Fig. 15.**
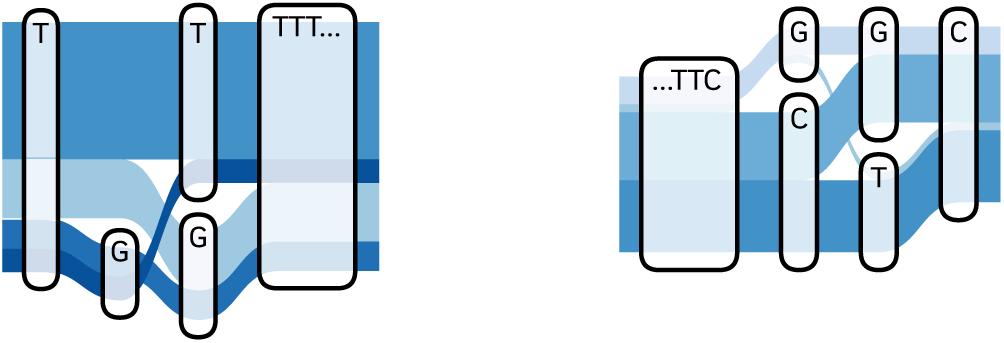
Two examples of abutting SNVs in the 1000 Genomes graph

**Fig. 16.**
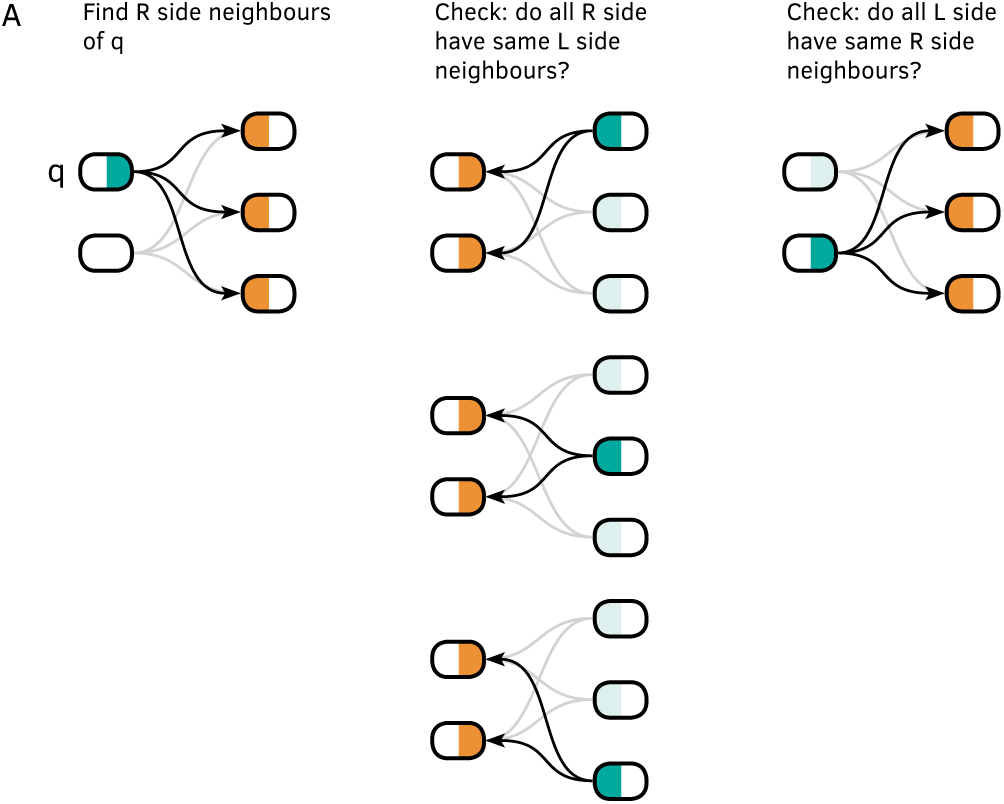
Illustration of Algorithm 1 returning a positive result

**Fig. 17.**
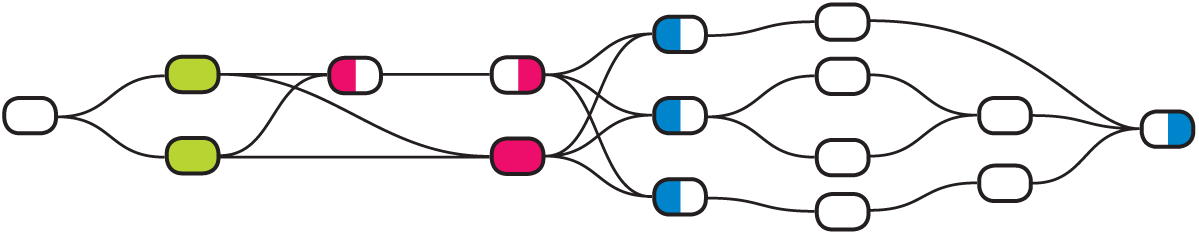
An ultrabubble decomposable into nested generalized sites; some sites marked

**Fig. 18.**
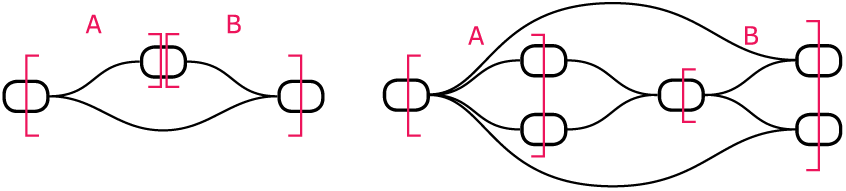
Two examples of deletion bundle-pairs

In order to prove that this is a valid algorithm for detection of balanced recombination bundles, we need the following lemma.

#### Lemma 23.

*Let* (*L, R*) *be a tuple of sets of node-sides*. *If* ∃*q ∈ L such that* ∀a *∈ Nb*(*q*), ∀b *∈ Nb*(*a*), *Nb*(*b*) ⊆⊆ *Nb*(*q*) *but Nb*(*q*) ⊂ *R, then* (*L, R*) *cannot be connected (in the sense of Definition 15)*.

*Proof. Let B* = ∪_*a*∈N*b*_(_*q*_) *Nb*(*a*). *We know that* ∀ *b ∈ B*, ∈ *Nb*(*b*) ⊆ *Nb*(*q*). *Suppose that* (*L,R*) *is connected. Choose r ∈ R*\*Nb*(*q*). *Then* ∃ *an internal chain c* = *q* → *r*_1_ → *ℓ*_1_ →· · · → *r_k_* → *ℓ_k_ →* r with r_i_ R, i_i_ *∈* L∀ i. Since q *∈* B, Nb(*b*) ∈ *Nb*(*q*) ∀ *b*∈ B, *and Nb*(*a*) ⊆ *B* ∈ *Nb*(*q*), *it is impossible that the sequence of node-sides c is both a valid internal chain and ends with r. Therefore* (*L, R*) *cannot be connected*.

#### Algorithm 1: Balanced recombination bundle finding

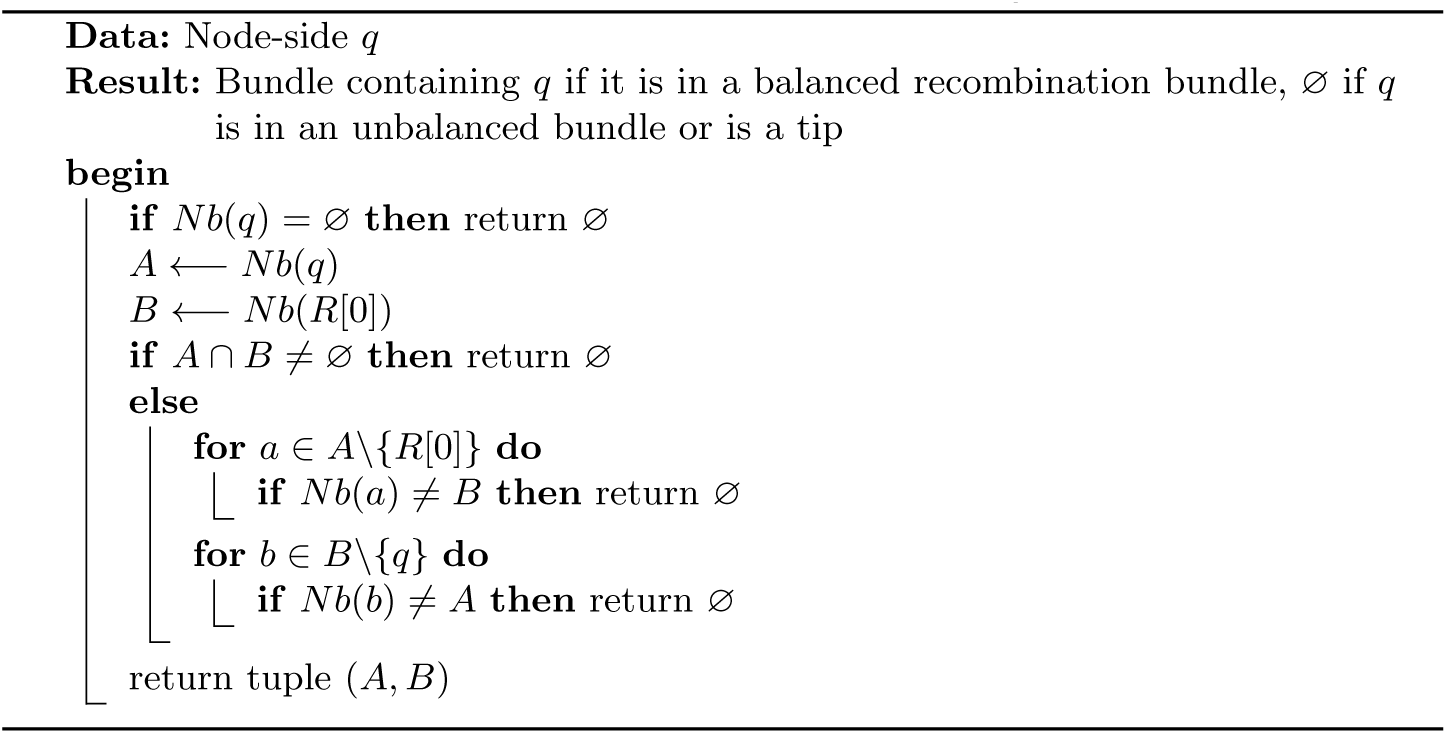

#### Proposition 24 (Validity of Algorithm 1).

*This algorithm detects all balanced recombination bundles, and rejects all unbalanced recombination bundles*.

*Proof. Suppose q is involved in a balanced recombination bundle* (*L, R*). *W.l.o.g. suppose that q* ∈ *L. Due to the complete matching property, the set Nb*(*q*) *in the algorithm is guaranteed to be equal to R. Due to the completeness property, the set Nb*(*R*[0]) *in the algorithm is guaranteed to be equal to L. It is evident that the algorithm directly verifies complete matching and acyclicity*.

*Suppose otherwise. Assuming we have eliminated all tips, which can be done in* 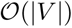 *time, Lemma 20 proves that q is involved in an unbalanced bundle B. If B fails acyclicity but not complete matching, then checking that A* ∩ *B* = ∅ *will correctly detect that L* ∩ *R* =∅.

*Otherwise, suppose that B fails complete matching. Suppose first that Nb*(*q*) ⊂ *R. We assert that* ∃*a* ∈ *Nb*(*q*) *such that* ∃*b* ∈ *Nb*(*a*) *such that* ∃*c* ∈ *Nb*(*b*) *such that* ∃c *Nb*(*q*). *This event will be detected by the second loop of the algorithm. This follows from the connectedness of B and Lemma 23*.

*Suppose otherwise that Nb*(*q*) = *R but* ∃*r* ∈ *R such that Nb*(*r*) ⊂ *L. Let c* ∈ *L\Nb*(*r*). *By matching*, ∃*r*′ ∈ *R such that r*′ ∈ *Nb*(*c*). *Therefore Nb*(*r*) *and Nb*(*r′*) *will be found to be unequal in the first loop of the algorithm*.

*Suppose otherwise that Nb*(*q*) = *R, Nb*(*r*) = *L*∀, *but* ∃ ℓ ∈L *such that Nb*(*ℓ*) ⊂R. *Then we will find in the second loop that Nb*(*ℓ*) ≠ *Nb*(*q*).

#### Proposition 25 (Speed of Algorithm 1).

*We can identify all balanced recombination bundles, all unbalanced bundles and all tips in* 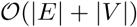 *time*.

*Proof. We depend on a neighbor index giving us* 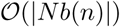 *iteration across neighbors of a node-side n*.

*We begin by looping over all node-sides and identifying all tips, which is achieved in* 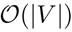 *time. We then loop again over all remaining node-sides. At each node-side q, we run the function describe above, which, if q is involved in a balanced recombination bundle, will return the bundle* 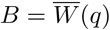. *It is evident that this function runs in* 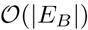 *time, seeing as it loops over each edge of B twice—once from each side—each time making an* 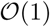 *set inclusion query. After B is built, all nodes are marked such that they are skipped when they are encountered in the global loop. This gives overall* 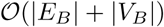 *exploration of B*.

*If q is involved in an unbalanced bundle* 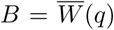, *this fact is detected by the same function in* 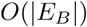 *time. In this case, we can find all nodes of B by performing a breadth-first search. Examination of the W-function will convince the reader that a breadth-first search will find all node-sides of B in* 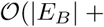 |*V_B_|) time. We follow the same procedure of marking all these node-sides to be skipped in the global loop*.

*This proves that, after eliminating tips in* 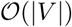time, we can build the set 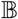 *of all non-isomorphic bundles B, and decide whether they are balanced recombination bundles, in time proportional to* 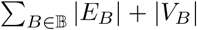 *But Lemma 20 and Corollary 22 tell us that* 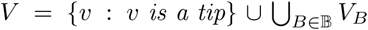. *and that all elements of this union of node-sides are disjoint. Furthermore, due to the matching property of bundles*, 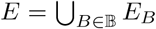, *and all elements of this union of edges are disjoint. Therefore, our method is* 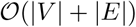.

### 5.3 Bundles and Snarl Boundaries

#### Definition 26.

*Given a “boundary” node-side b* = *s or t of a snarl* (*S, s, t*), *we call the tuple* (*b, Nb*(*b*)) *a snarl comb. A snarl comb is called proper if* ∀*n* ∈ *Nb*(*b*), *Nb*(*n*) = {*b*} *and b* ∉ *Nb*(*b*).

It is easy to verify that a *proper snarl comb* is a balanced recombination bundle. It is also easy to see that an improper snarl comb is, according to set inclusion of tuples, a proper subset of a unique bundle.

#### Proposition 27 (Bundles do not cross snarl boundaries).

*Let* (*S, s, t*) *be a snarl. Suppose that B* = (*L, R*) *is a bundle. Then either all node-sides involved in B are members of S, or no node-side involved in B is a member of S*.

*Proof. Suppose that there exists a bundle B* = (*L, R*) *with node-sides both within S and not within S. Let x,y be involved in B, with x ∈S, y* ∉ *S. W.l.o.g., suppose x* ∈ *L, y* ∈ *R. This implies that there exists an internal chain p* = *x* →· · · → *y*. *But then this implies that there exists a* ∈ *S, b* ∉ *S such that a* ∈ *Nb*(*b*), *which would allow us to use the edge a* →*b to create a path violating the 2-node separability of S*

### 5.4 Defining Sites using Bundles

#### Definition 28.

*An ordered pair* (*B*_1_, *B*_2_) *of balanced recombination bundles is compatible if either*

1. 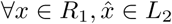 and 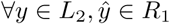
2. ∃ *a bijection f* : *L*_1_ → *R*_2_ *such that* ∀*x* ∈*R*_1_, *there exists a unique path p*(*x*) *from x*→ · · · → f (*x*), *and all paths p*(*x*) *are disjoint*.

#### Definition 29.

*If two recombination bundles are compatible, we define the set p*(*x*) *to be a bundled simple site P*.

*Claim*. Consider a bundled simple site *P* in a graph G, lying between compatible balanced recombination bundles *B*_1_, *B*_2_. The set of all paths in *G* which contain paths *p* ∈*P* as contiguous subpaths is isomorphic to the set-theoretic product *P*(*L*_1_) × *P* × *P*(*R*_2_) consisting of the three sets

1. *P*(*L*_1_) := {paths in *G\S* terminating in *x*, for some *x*∈ *L*_1_}
2. *P*
3. *P*(R_2_) := {paths in *G\S* beginning with *y*, for some *y* ∈ *R*_2_}
 under the function mapping *p_1_* ∈*P|*(*L_1_*),*p* ∈ *P, p_2_* ∈ *P*(R_2_) to their concatenation.

We will call a balanced recombination bundle *B* = (*L, R*) *trivial* if both *L* and *R* are singleton sets.

#### Definition 30.

*An ultrabubble* (*U, s, t*) *is a generalized simple bubble if*

1. ({*s*}, *Nb*(*s*)) *and* (*Nb*(*t*), {*t*}) *are balanced recombination bundles*
2. *The set of all non-trivial balanced recombination bundles admits a linear ordering X* → *B*_1_ → … *B_k_* → *Y such that X and Y are either of* ({*s*}, *Nb*(*s*)) *and* (*Nb*(*t*), {*t*}), *X is compatible with B*_1_, *every B*_*i*_ *is compatible with B_i_*_+1_, *and B_k_ is compatible with Y*

#### Definition 31.

*An ultrabubble U is decomposable into nested generalized sites if either:*

1. *It is a generalized simple bubble*
2. *When each ultrabubble* (*V, u, υ*) *nested in U which is a decomposable into nested generalized sites is replaced with a single edge spanning u and υ, then U is a generalized simple bubble*

We sketch a linear-time method of building sites from a tree diagram of nested ultrabubbles. We run Algorithms 2 and 3 starting at bottom-level nested ultrabubbles. If ultrabubble has all nontrivial balanced recombination bundles paired, then, when we evaluate the ultrabubble containing it, we represent it as a single edge from its source to sink.

In Algorithm 3, which follows below, we refer to the individual sets of node-sides forming the tuples (*L, R*) of a bundle as bundle-sides.

### 5.5 Bundles Containing Deletions

Our bundles—and therefore our sites—fail to detect the graph motifs formed by deletions spanning otherwise well-behaved variants. We define a special, well-behaved subclass of unbalanced bundle to address this.

#### Definition 32.

*A deletion bundle-pair is a tuple* (*L_A_, R_A_, L_B_, R_B_*) *such that*

1. ∀ℓ ∈ *L_A_*, ∀ _*r*_ ∈ *R_A_*, {*ℓ, r*} ∈ *E*
2. ∀ ℓ∈ *L_A_*, ∀_*r*_ ∈ *R_B_*, {*ℓ, r*} ∈ *E*
3. ∀ ℓ∈ *L_B_*, ∀_*r*_ ∈ *R_B_*, {*ℓ, r*} ∈ *E*
4. ∃ *no other edge involving any node-side n* ∈ *L_A_, L_B_, R_A_ or R_B_*

#### Algorithm 2: Finding spans connecting bundles

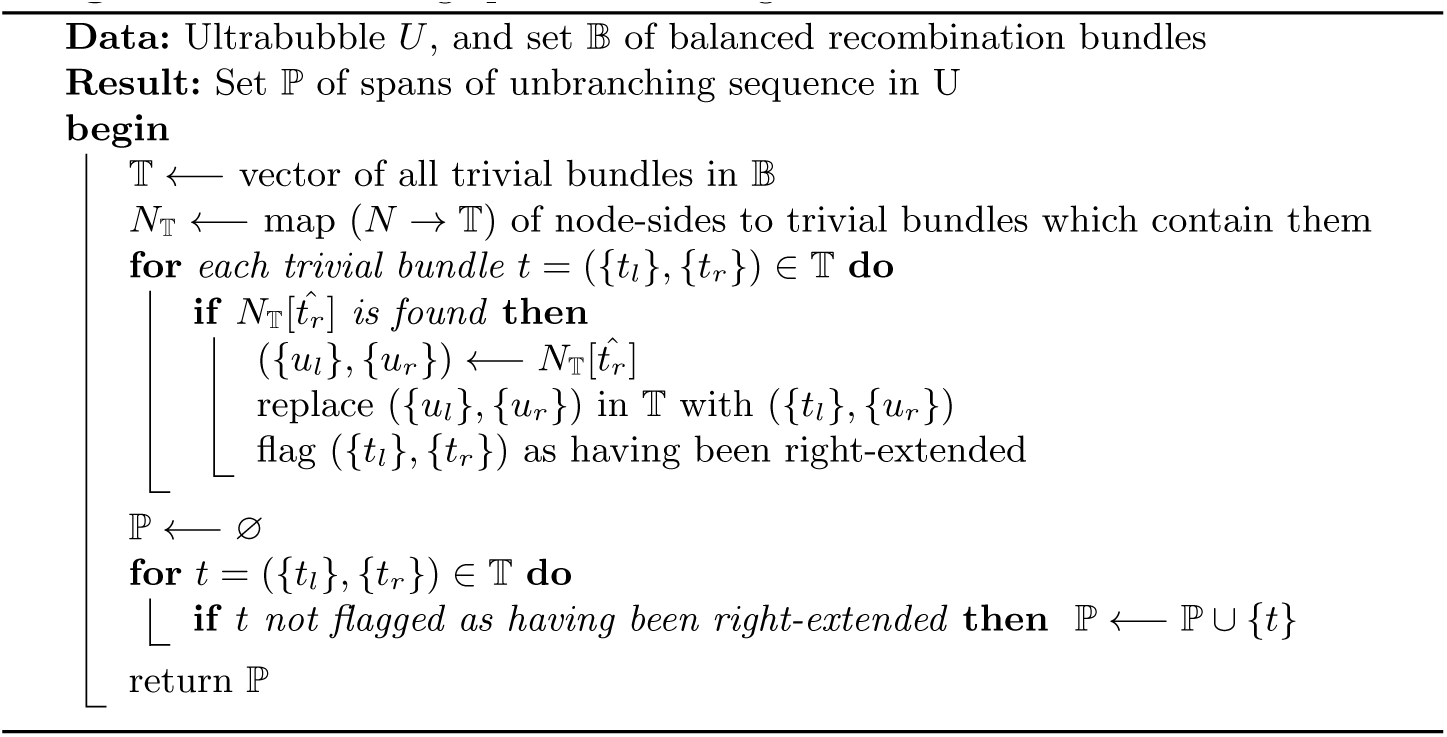

#### Algorithm 3: Finding compatible bundles

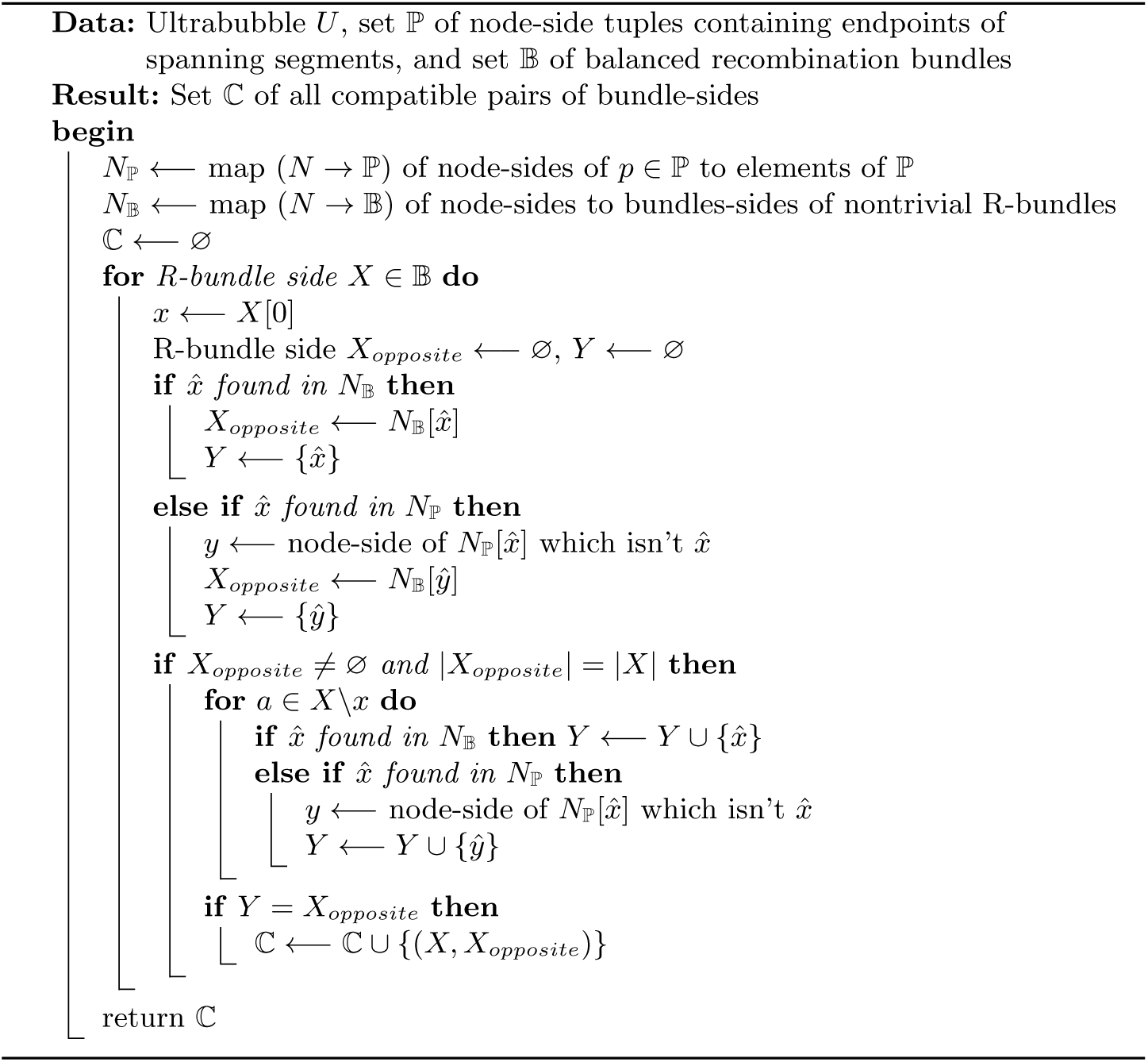

These structures occur when two balanced recombination bundles on either side of some span of graph are bridged by deletions. It remains necessary to check that there is graph structure joining the nodes of *R_A_* to *L_B_* for this to be the case.

Algorithm 4 below will detect deletion bundle pairs from among the set of unbalanced bundles in linear time.

#### Proposition 33.

*Given a set of acyclic unbalanced bundles, this algorithm finds those among then which are deletion bundle pairs*.

*Proof. Suppose that q is involved in a deletion bundle pair* (*L_A_, R_A_, L_B_, R_B_*). *W.l.o.g, either q* ∈ *L_A_ or q* ∈ *L_B_*.

*Suppose first that q* ∈ *L_B_* : *In this case, Nb*(*q*) = *R_B_*. *We then know that* ∀*a ∈ RB, Nb*(*a*) = *LA∪LB. This will trigger the condition L*_2_ = ∅. *The elements of a ∈ L*_1_ *will segregate into precisely two groups: one such that Nb*(*a*) = *R_B_* — *the elements a ∈ L_B_, and another group such that Nb*(*a*) = *RA* ∪ *RB* — *the elements a* ∈ *L_A_*. *If these conditions are fulfilled, we then build R_A_ and R_B_. It remains to verify that* ∀*b ∈ R_A_, Nb*(*b*) = *L_A_, and* ∀ _b_ *∈ R_B_, Nb*(*b*) = *L_A_ ∪ L_B_*.

*Suppose otherwise that q G L_A_*: *In this case, Nb*(*q*) = *R_A_* ∪ *R_B_*. *This will trigger the condition L*_2_ ≠∅ *since the elements b* ∈ *Nb*(*q*) *will segregate into two groups: R_A_, where if b G R_A_, Nb*(*b*) = *L_A_ and R_B_, where if b* ∈ *R_A_, Nb*(*b*) = *L_A_* ∪ *L_B_*. *If this condition is met, then it remains to check that* ∀*a* ∈ *L_A_, Nb*(*a*) = *R_A_* ∪ *R_B_* and ∀*a* ∈ *L_B_, Nb*(*a*) = *R_B_*.

*Suppose otherwise that q is not involved in a deletion bundle pair. Suppose that Algorithm 4 does not fail, returning. ∅ There are two possibilities then for the nature of the unbalanced bundle* (*L, R*) *for which q* ∈ *L*.

*First, suppose the condition L*_2_ = ∅ *was triggered. The* ∃*q* ∈ *L such that, where L* ′ := {*l* ∈ *L* |*l* ∈ *Nb*(*a*) *for some a* ∈ *Nb*(*q*)}, *Nb*(ℓ) *C Nb*(*q*) ∀ ℓ∈ *L* ′. *Then by Lemma 23, Nb*(ℓ)≥ *Nb*(*q*)∀ℓ∈ *L. Therefore it must be that Nb*(*q*) = *R. Furthermore, to pass the search for R_A_, there must* ∃ *R_A_ such that if* ℓ∈ *L and Nb*(ℓ) ≠ *R, then Nb*(ℓ) = *R_A_. Furthermore, to pass the conditions of the subsequent two loops, it must be that* ∀*r* ∈ *R\RA, all Nb*(*r*) *are the same, and* ∀*r* ′∈ *R_A_, all Nb*(*r*′) *are the same. Furthermore, to pass the last condition checked, must be that Nb*(*r*′>) *from the latter group* ⊂*Nb*(*r*). *And since L_A_* :={ℓ∈ *L* | *Nb*(ℓ) = *R*} *and L_B_* := {*ℓ∈ L* | *Nb*(ℓ) = *R_B_*} *are such that L_A_ ∩L_B_* = ∅ *L_A_* ∪ *L_B_* = *L, these conditions all together ensure that* (*L, R*) *is a deletion bundle pair*.

*Otherwise, Nb*(*q*) *segregates into two disjoint subsets R_A_* := {*r* ∈ *Nb*(*q*)| *Nb*(*r*) = *L_A_*}, *R_B_* := {*r* ∈ *Nb*(*q*) | *Nb*(*r*) = *L_A_ ∪L_B_ for some L_A_, L_B_ ⊂L such that L_A_ ∩ L_B_* = ∅. *To pass further conditions, it is necessary that ∀ ℓ∈ L_B_, Nb*(*l*) = *R_B_ and* ∀*ℓ* ∈ *LA, Nb*(ℓ) = *R_A_*∪ *R_B_*. *It remains to show that LA ∪L_B_* = *L and R_A_ ∪ R_B_* = *R, these can be proven by application of Lemma 23. Therefore in this case, it must also be that* (*L, R*) *is a deletion bundle pair*.

#### Algorithm 4: Deletion bundle pair finding

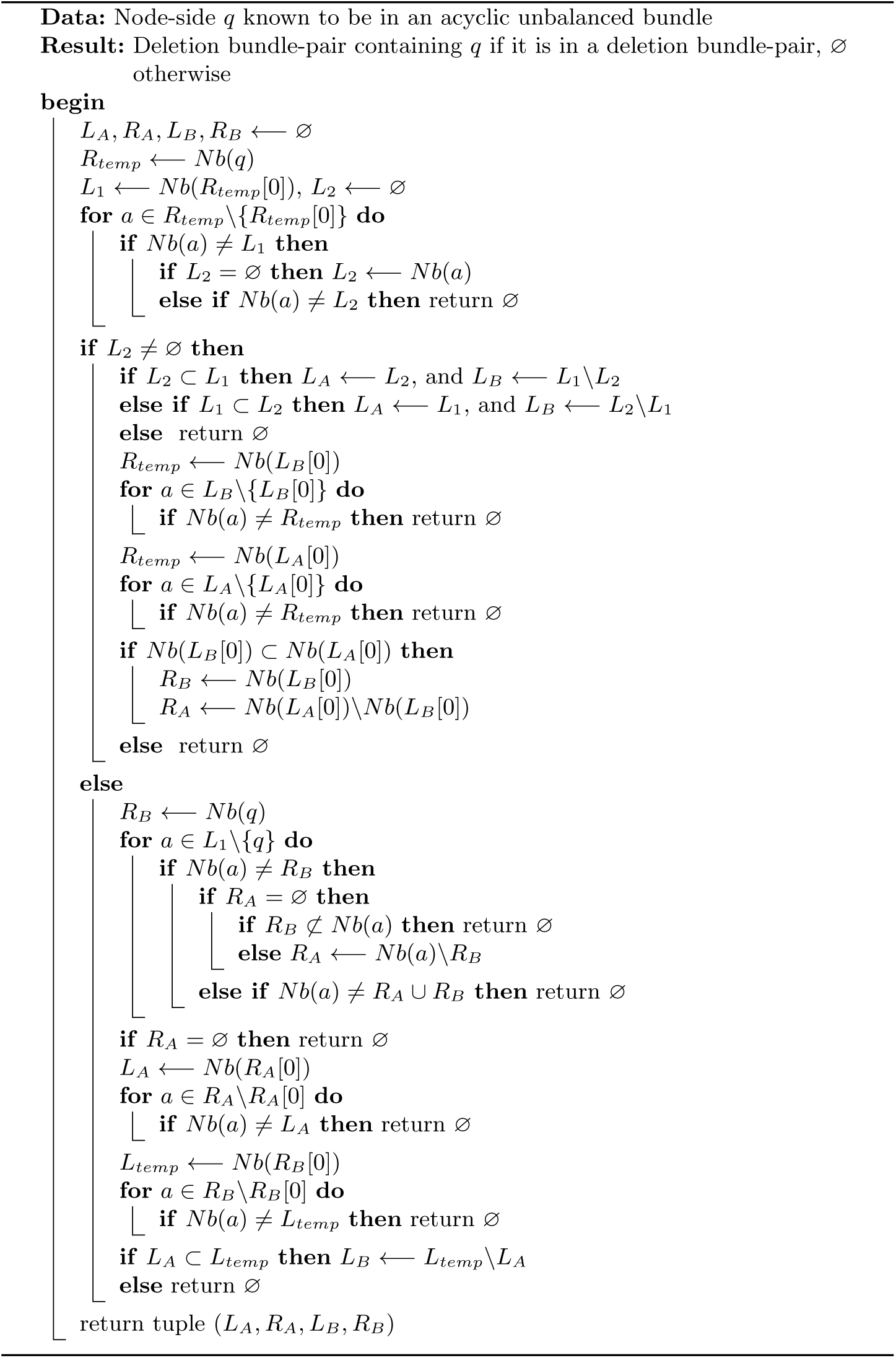

#### Proposition 34.

*This algorithm finds deletion bundles in 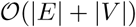 time*.

*Proof. Note that a deletion bundle-pair is a special type of unbalanced bundle. Therefore, if, given an unbalanced bundle B, we can check whether it is a deletion bundle-pair in 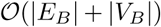 time, by the arguments of Proposition 21, we can find all deletion bundle-pairs in 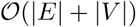time*.

*Inspection of the algorithm shows that, like the algorithm for identifying balanced recombination bundles, it performs two 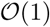 set-inclusion queries per edge, making it 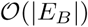 overall*.

## Discussion

Graph formalism has the potential to revolutionize the discourse on genetic variations by creating a model and lexicon that more fully embraces the complexity of sequence change. This is vital: the current linear genome model of a reference sequence interval and alternates is insufficient. It fails to express nested variation and can not properly describe information about the breakpoints that comprise structural variations.

The introduction, in order, of bubbles, superbubbles, ultrabubbles and snarls progressively generalizes the concept of a genetic site to accommodate more general types of variation using progressively more general graph types. In this paper we both review and build on these developments, showing how the recently introduced ultrabubbles can be furthered sub-classified using concepts from circuit theory. This expands the simple notion of proper nesting described in the original ultrabubble paper. Furthermore, we describe how we can extend the theory of ultrabubbles by generalizing ultrabubble boundaries to another sort of boundary structure—the bundle—which allows us to describe regions where variants are packed too closely to be segregated into separate ultrabubbles.

Our methods are powerful in decomposing dense collections of nested or closely packed variation into meaningful genetic sites. We anticipate that these structures will become increasingly common in the analysis of variation using graph methods, as sequencing datasets containing variation from increasing numbers of individuals become available.

## Acknowledgements

Y.R. is supported by a Howard Hughes Medical Institute Medical Research Fellowship. This work was also supported by the National Human Genome Research Institute of the National Institutes of Health under Award Number 5U54HG007990 and grants from the W.M. Keck foundation and the Simons Foundation. The content is solely the responsibility of the authors and does not necessarily represent the official views of the National Institutes of Health. We thank Wolfgang Beyer for his visualizations of 1000 Genomes data in a variation.

## Footnotes

It is on bidirected graphs

